# Resting-state fMRI signals contain spectral signatures of local hemodynamic response timing

**DOI:** 10.1101/2023.01.25.525528

**Authors:** Sydney M. Bailes, Daniel E. P. Gomez, Beverly Setzer, Laura D. Lewis

## Abstract

Functional magnetic resonance imaging (fMRI) has proven to be a powerful tool for noninvasively measuring human brain activity; yet, thus far, fMRI has been relatively limited in its temporal resolution. A key challenge is understanding the relationship between neural activity and the blood-oxygenation-level-dependent (BOLD) signal obtained from fMRI, generally modeled by the hemodynamic response function (HRF). The timing of the HRF varies across the brain and individuals, confounding our ability to make inferences about the timing of the underlying neural processes. Here we show that resting-state fMRI signals contain information about HRF temporal dynamics that can be leveraged to understand and characterize variations in HRF timing across both cortical and subcortical regions. We found that the frequency spectrum of resting-state fMRI signals significantly differs between voxels with fast versus slow HRFs in human visual cortex. These spectral differences extended to subcortex as well, revealing significantly faster hemodynamic timing in the lateral geniculate nucleus of the thalamus. Ultimately, our results demonstrate that the temporal properties of the HRF impact the spectral content of resting-state fMRI signals and enable voxel-wise characterization of relative hemodynamic response timing. Furthermore, our results show that caution should be used in studies of resting-state fMRI spectral properties, as differences can arise from purely vascular origins. This finding provides new insight into the temporal properties of fMRI signals across voxels, which is crucial for accurate fMRI analyses, and enhances the ability of fast fMRI to identify and track fast neural dynamics.

## Introduction

Functional magnetic resonance imaging (fMRI) enables non-invasive measurement of human brain activity via the hemodynamic response. When activity in a population of neurons changes, these changes give rise to the blood-oxygenation-level-dependent (BOLD) signal measured in most fMRI studies (1). Thus far, however, BOLD fMRI has exhibited relatively limited ability to provide the fine-grained temporal information necessary for deepening our understanding of brain dynamics. This is due to the fact that the signals obtained from BOLD fMRI are not direct measures of neural activity, but rather reflect the coupling between neuronal activity and the hemodynamic response, which evolves on a time course of seconds (2, 3). This coupling between neural activity and the BOLD signal can be represented by the hemodynamic response function (HRF) (4, 5). The properties of the HRF depend on many interconnected factors, including the effects of local vascular architecture and cerebrovascular dynamics, that vary substantially across the brain and between individuals (4–7). The relative timing and shape of the HRF, therefore, also varies considerably across brain regions and even between neighboring voxels (8–13). Hemodynamic response temporal lag variation is substantially larger than many neural effects of interest, introducing variability on the order of several seconds (8–11). Thus, to enable inferences about the relative timing of neural activity using signals obtained from BOLD fMRI, it is crucial to understand the variations in HRF timing across the brain.

Advances in acquisition technology now allow high-resolution whole brain fMRI data to be acquired at fast (<500 ms) rates (14–19), suggesting that fMRI could provide a unique tool to noninvasively track temporal sequences of neural activity (20) across the entire brain. Indeed, recent studies have revealed highly structured temporal dynamics using fMRI and suggest that fMRI can enable whole-brain mapping of temporal sequences (21–25). Furthermore, hemodynamic signals have been shown to contain more information about fast and high-frequency activity than previously thought (21, 26–29). Studies examining individual brain regions have demonstrated that fMRI can achieve impressive temporal precision within regions, on the order of 100 ms (30), meaning that high fidelity temporal information is present within these hemodynamic signals. However, a key remaining challenge is that the hemodynamic differences across the brain confound our ability to infer the timing of the underlying neural activity from BOLD fMRI. Specifically, if a given brain region shows earlier BOLD activity, it could be due to faster neural activity or simply due to a faster hemodynamic response in that region. Fully exploiting the higher temporal resolution provided by fast fMRI techniques will therefore ultimately require accounting for differences in the temporal dynamics of the hemodynamic response across the whole brain.

Despite the well-known heterogeneity of hemodynamic timing, most common analysis approaches for BOLD fMRI data assume a standard, canonical HRF shape throughout the brain (31). This approach is understandable, since the true HRF is not known, but it nevertheless cannot account for the vascular confound introduced by hemodynamic response variability. Incorrect assumptions about the shape and timing of the HRF can lead to incorrect inferences regarding the underlying neural activity (32–34), and studies that assume a whole brain canonical HRF are unable to decouple the neural and vascular components of the BOLD signal (5, 35–38). Even when using flexible modeling approaches, such as basis sets or finite impulse response models, it is not possible to determine whether a given region’s faster fMRI response reflects fast neural activity, or simply faster local neurovascular coupling (34).

The fact that most studies do not account for variations in HRF dynamics is largely due to methodological challenges. Previous work has demonstrated that it is possible to quantify hemodynamic lags across brain regions, and even on a voxel-wise level, to detect the relative order of BOLD responses with high temporal precision (11–13, 30, 38–47). One such method is the use of a stimulus paradigm that drives activity in particular brain regions where the neuronal response properties are relatively well understood and controlled such as primary sensory or motor cortices (5, 10–12, 40). However, this approach cannot be applied to the majority of the brain, where the neuronal response properties are not known ahead of time. Alternatively, using a breath hold or similar hypercapnic challenge can modulate cerebral blood flow (CBF) to all vascularized regions with minimal changes in cerebral metabolic rate of oxygen (CMRO_2_), allowing mapping of vascular latencies (38, 39, 42–45, 48). However, breath hold tasks are not suitable for all subject populations, as some patients may have difficulty complying with the breath hold task. Furthermore, breath hold tasks modulate and measure cerebrovascular reactivity (CVR), which contributes to neurovascular coupling but is a distinct process. While neurovascular coupling reflects the alterations in local hemodynamics that occur in response to changes in neural activity, CVR is specifically a measure of a blood vessel’s capacity to dilate and constrict in response to a vasoactive stimulus, and does not include the extensive metabolic and molecular factors that also drive neurovascular coupling (48–50). In fact, there is evidence that while CVR is affected by healthy aging, some metrics of neurovascular coupling are not, hinting that distinct mechanisms may shape these two patterns (51–53).

A task-free, neurovascular-based approach for detecting the lags of intrinsic neurovascular coupling would therefore be broadly relevant for analyzing fMRI data. A potential alternative route towards identifying local hemodynamic properties is to examine the properties of resting-state fMRI data. Resting-state fMRI signals reflect neurovascular coupling induced by spontaneous neural activity (54, 55) and confer the additional benefit of being task-independent, which makes it a viable scan type for patient populations. Moreover, unlike stimulus- or task-based paradigms for mapping local hemodynamic response timings in particular brain regions, resting-state fMRI can be used to examine the HRF across the whole brain. These advantages have prompted past research into the utility of resting-state fMRI signals to estimate the HRF itself with prior work utilizing deconvolution approaches to explore HRF timing in resting-state data (46, 47, 56). However, these approaches require assumptions about the underlying neural events, which are not known. We therefore investigated whether intrinsic signatures of local neurovascular coupling dynamics are present in the resting-state fMRI signal.

Our goal was to understand whether information about the temporal dynamics of the hemodynamic response is present in resting-state fMRI data. We first used simulations of the BOLD response to illustrate how distinct, physiologically relevant HRF shapes should produce marked differences in the frequency content of resting-state signals. Next, we verified this result in fast fMRI data collected at 7 Tesla using visual stimulation to induce a neural response with known timing in primary visual cortex. We quantified the temporal delay of voxels in the primary visual cortex (V1) in response to a controlled, oscillating visual stimulus, and found that voxels with fast and slow hemodynamic responses exhibited distinct resting-state spectral features. We further extended our analyses to the visual thalamus (lateral geniculate nucleus, LGN) and found that this principle generalized to subcortex. To understand the potential of this information as a tool to predict the temporal properties of individual voxels, we then trained classifiers to use information from the resting-state spectrum to classify voxels as being fast or slow cortical voxels, or even faster LGN voxels. We found that resting-state signals were better predictors of voxel-wise differences in relative hemodynamic timing than latencies measured from a gold standard breath hold task. Our results establish that information about hemodynamic timing can be extracted from the frequency spectrum of resting-state fMRI signals. This demonstrates that resting-state fMRI can provide a way to understand and predict the temporal dynamics of the HRF across the brain, which is critical for interpreting neural activity using BOLD fMRI.

## Results

### The temporal dynamics of the HRF profoundly impact the spectrum of simulated BOLD responses

Previous modeling work has illustrated that narrower HRFs should result in BOLD responses containing more high frequency power (27), suggesting that local variations in HRF timing should manifest as local variations in the frequency content of BOLD signals. We therefore hypothesized that the frequency spectrum of BOLD dynamics in the resting-state can be used to infer the relative timing of the task-driven hemodynamic response. We first aimed to illustrate this property by simulating the BOLD frequency response using HRFs with different temporal dynamics. If we assume that the BOLD response is a linear time-invariant system, we can compute the BOLD response as a convolution between a given input (i.e., the stimulus) and the characteristic input response of the system (i.e., the HRF). Then, by varying the frequency of the stimulus we can construct a spectrum of the BOLD frequency response for HRFs with faster or slower dynamics (Fig. 1A). We performed this simulation using six different HRFs (Fig. 1B) with physiologically representative values for their time-to-peak (TTP), full width at half maximum (FWHM), and amplitude (11). We observed that while HRFs with faster dynamics produced less power in the low frequency bands compared to those with slower dynamics, they also showed a shallower decline in power at higher frequencies (Fig. 1C). Furthermore, this effect was preserved when we normalized the different HRFs to have the same peak amplitude (Supplementary Fig. S1A), demonstrating that this phenomenon is not solely due to the higher amplitude of slower HRFs (Supplementary Fig. S1B). This effect was also preserved if we accounted for the 1/f-like spectral pattern that neural activity displays (Supplementary Fig. S1C-E). These simulations demonstrate that there should be profound differences in the relative power at low versus high frequencies for voxels with fast vs. slow HRFs which we hypothesize can be quantified and related to the temporal dynamics of the hemodynamic response. This relationship will be explored in the remainder of the paper.

**Figure 1.**
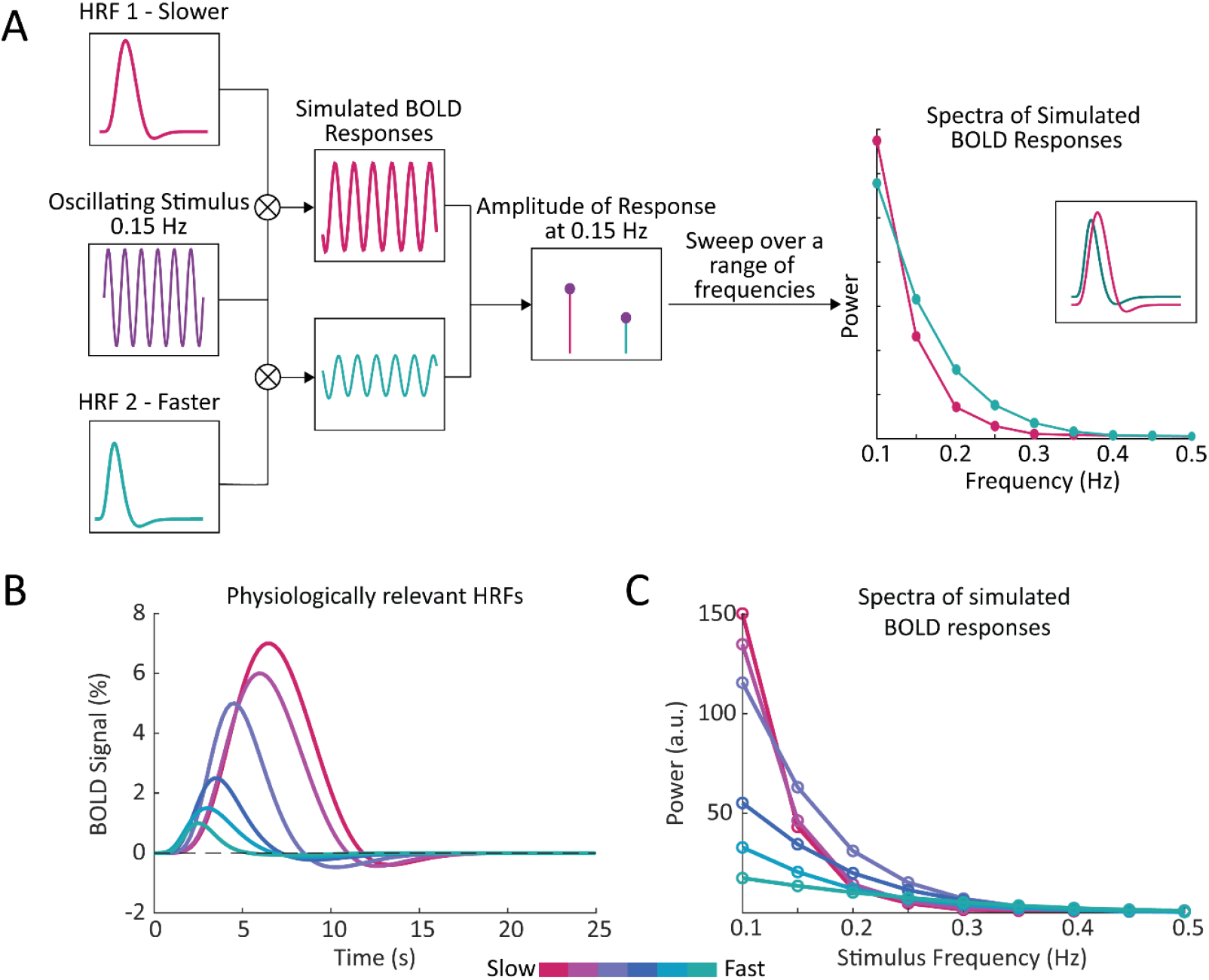
Simulations show that the temporal properties of the hemodynamic response function affect the frequency spectrum of the BOLD signal. **A)** We generated a simulated BOLD response to determine the response amplitude of each HRF to each neural frequency. By convolving a given HRF with an oscillating stimulus, and sweeping across a range of frequencies, we generated a frequency spectrum of the simulated BOLD responses. This simulation was repeated using HRFs with varying temporal properties, to compare the frequency spectrum of simulated BOLD responses with faster or slower hemodynamic responses. **B)** We generated a range of HRFs with physiologically plausible timings and amplitudes (*11*). **C)** We found that temporal properties of the HRF had noticeable effects on the simulated spectra, particularly under 0.2 Hz.

### Features of the resting-state spectrum show significant differences between fast and slow voxels

Based on these simulation results, we then examined the frequency content of resting-state fMRI data. Spontaneous BOLD fluctuations captured in resting-state fMRI are linked to fluctuations in neuronal activity (54, 55, 57), and accordingly, reflect the neurovascular coupling mechanisms that link neural fluctuations to BOLD fluctuations. To empirically test the prediction that fast and slow voxels should have distinct frequency content in the resting-state, we first used a task paradigm to identify voxels in the primary visual cortex (V1) with consistently fast or slow hemodynamic responses. To drive continuous oscillations in V1 we presented the subjects with a 12-Hz counterphase flickering radial checkerboard with the luminance contrast of the checkerboard modulated in time as a sine wave of 0.05 Hz (Fig. 2A, top). We used a combination of an anatomical and functional localizer to identify stimulus-driven voxels in V1 (Fig. 2B). For those voxels that were significantly driven by the stimulus, we then calculated the phase lag, or relative response latency to the stimulus (Fig. 2C). Consistent with prior studies (10, 28, 58) we found a wide range of hemodynamic response lags within V1 (Fig. 2A, bottom). We extracted groups of fast and slow responding voxels (Fig. 2D-E). Then, for each voxel identified as fast or slow within the task run, we calculated that voxel’s frequency spectrum in the resting-state. Figure 2F shows the resting-state spectrum of a representative fast and slow voxel from a single subject where the differences in the spectra, particularly the distinct slope under 0.2 Hz, are visible.

**Figure 2.**
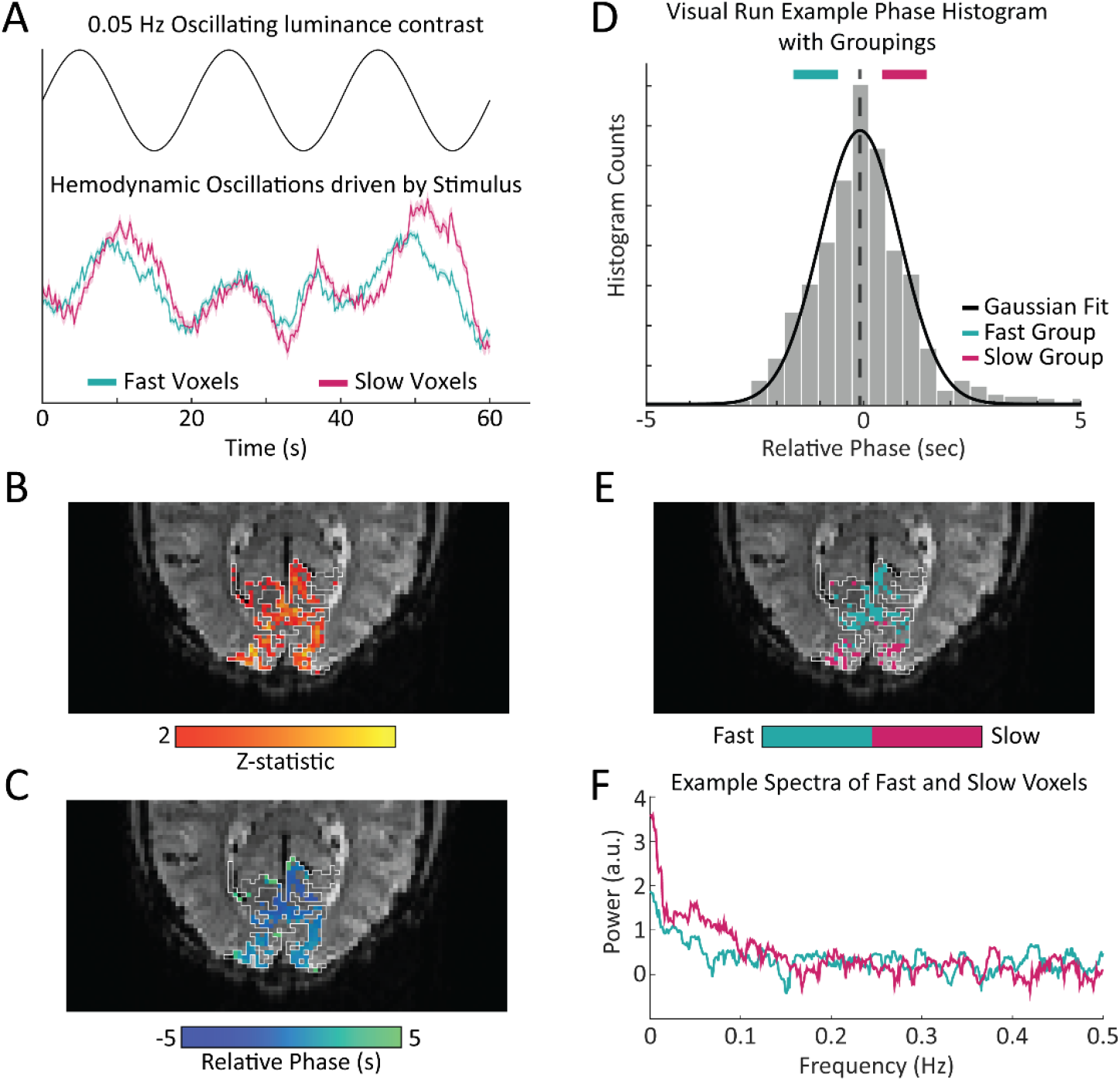
Experimental design: oscillating visual stimuli identify fast- and slow-responding voxels in V1. **A)** Subjects viewed a flickering checkerboard with oscillating luminance contrast to drive neural oscillations in V1. Some voxels showed a faster response to the visual stimulus and other showed a slower response, with a noticeable difference in the temporal dynamics of the mean response in these groups. Shading represents standard error. **B)** Example of a functional localizer in one subject with the white lines denoting the outline of the primary visual cortex (V1) based on anatomical segmentation. One visual stimulus run was used as a functional localizer to identify stimulus-driven voxels in V1. **C)** For all stimulus-driven voxels in V1, the phase of the response to the visual stimulus was calculated from the average of the visual stimulus runs not used as the functional localizer, corresponding to the local hemodynamic delay. **D)** We defined groups of “fast” and “slow” voxels using a Gaussian fit to the histogram of phases. Histogram shows example from one representative subject. **E)** Example map of fast and slow voxels generated for a single subject. **F)** Frequency spectrum of a representative slow and fast voxel’s resting-state signal, showing a difference in power drop off across frequencies, with a steeper slope for the slower voxel.

Our simulations had predicted a difference in the overall frequency content of resting-state fMRI signals in voxels with fast versus slow HRFs. To quantify this property across voxels, we sought to generate a set of spectral features that could capture these resting-state spectral dynamics. We constructed four spectral features to capture spectral properties: the slope using a linear fit under 0.2 Hz, the exponent of an aperiodic 1/f fit, the amplitude of low frequency fluctuations (0.01-0.1 Hz power; ALFF) (59), and the fractional ALFF (ratio of 0.01-0.08 Hz to 0-0.25 Hz; fALFF) (60)(Fig. 3). Each feature of the resting-state spectra revealed significant differences (Wilcoxon rank-sum test, p<0.05) between fast and slow voxels across subjects. The slope showed significant differences between the fast and slow voxels within each individual subject (15/15), while the aperiodic exponent, ALFF, and fALFF showed significant differences in 14, 11, and 13 subjects, respectively. (See Supplementary Table S1 for p-values). Each of these features reflects information about the frequency content at high and low frequencies, suggesting that this was an effective metric for differentiating voxels with fast or slow hemodynamic responses. Notably, the most effective features for distinguishing fast and slow voxels were the ones that explicitly captured the relative difference in high-frequency vs. low-frequency power.

**Figure 3.**
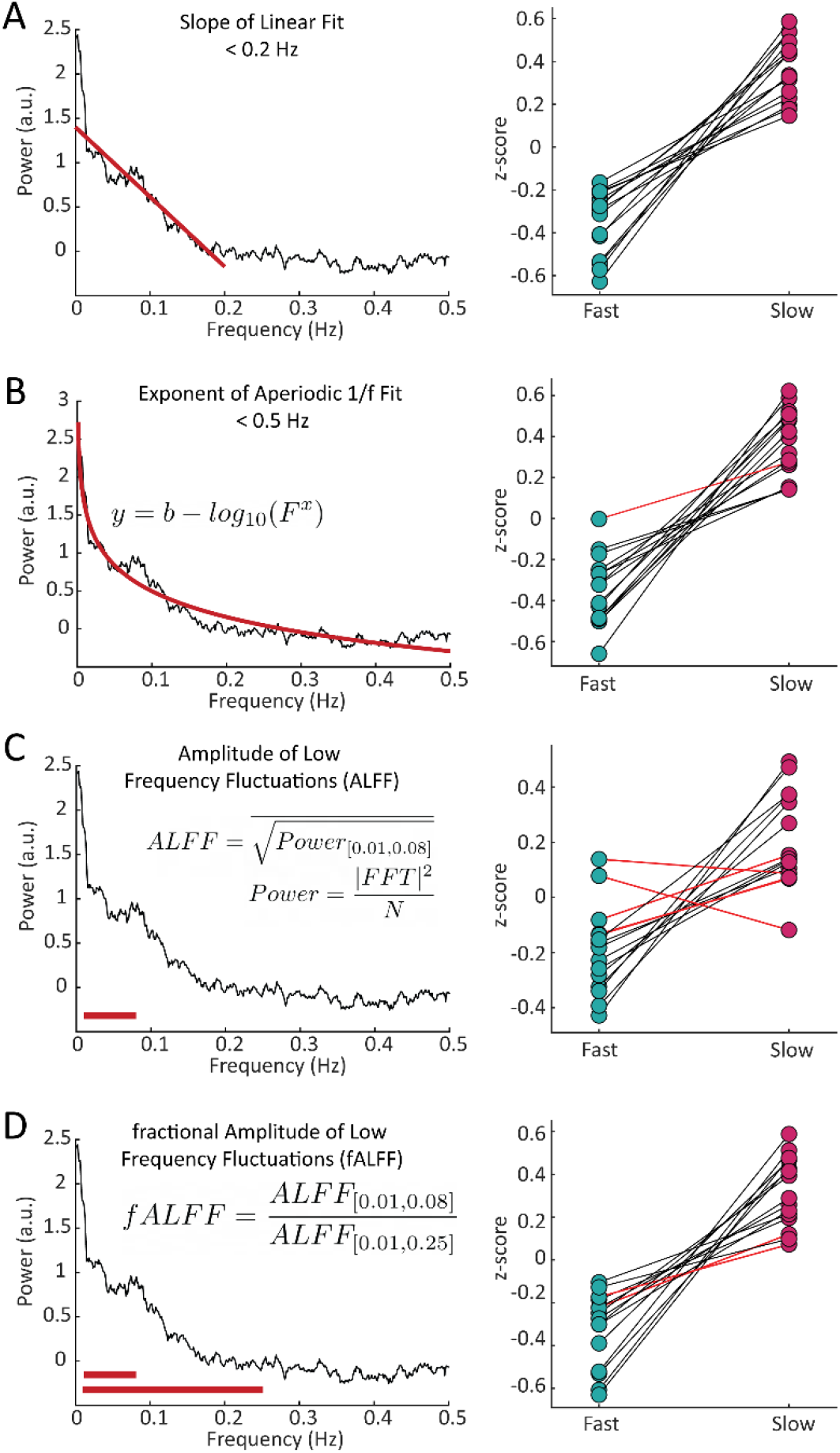
Features of the resting-state spectrum differed between fast and slow voxels in each subject. For each subject, we calculated four features of the resting-state frequency spectrum and compared the values between the task-defined fast and slow voxels using a Wilcoxon rank sum test. For the **A)** slope, using a linear fit of frequency spectrum under 0.2 Hz, 15/15 subjects showed significant differences; **B)** exponent of an aperiodic 1/f fit under 0.5 Hz, 14/15 subjects showed significant differences; **C)** amplitude of low frequency fluctuations (ALFF), 11/15 subjects showed significant differences; and **D)** fractional ALFF, 13/15 subjects showed significant differences. Black lines indicate a significant difference in a given subject (Wilcoxon rank-sum test, p < 0.05) and red lines indicate a non-significant difference.

### Faster hemodynamic responses in thalamus are also reflected in shallower frequency spectra

A key strength of fMRI is its ability to noninvasively image activity in the subcortex. Having established that the resting-state spectrum contained signatures of local hemodynamic response timing within V1, we next aimed to test whether this principle extended to the visual thalamus, specifically the LGN. Prior studies have shown that LGN has faster hemodynamic responses than V1 (61–63); however, due to its small size and lower signal-to-noise ratio, extracting its spectral features accurately could be more challenging. We generated individual masks of the LGN within each subject using the individual anatomical segmentation (64) and a functional localizer (Fig. 4A). To confirm the presence of stimulus-locked oscillations in the LGN and to assess the relative timing of its response, we first examined the latency of the average response to the visual stimulus in the LGN, as compared to the fast and slow groups in V1. We observed that the LGN peaked before both the fast and slow groups in V1 (Fig. 4B), consistent with prior work demonstrating faster hemodynamics in thalamus (61–63). Then, to test whether this faster hemodynamic response was similarly linked to flatter resting-state spectra, we compared the LGN voxels’ resting-state features to the previously identified fast and slow voxels in the cortex. We found that for all subjects there were clear differences in each spectral feature within the LGN compared to both the fast and slow voxels of the visual cortex (Fig. 4C-F). (See Supplementary Table S1 for p-values). The feature that had the poorest sensitivity to differences between the LGN features and cortical features is ALFF, which could be explained by ALFF’s higher sensitivity to non-neural noise sources (60). We thus observed even shallower frequency slopes for the fast LGN voxels – again consistent with our simulation results, demonstrating that this pattern held not just within V1 but even extended to the LGN of the thalamus. This observation was also robust to controlling for the higher thermal noise in LGN signals (Supplementary Fig. S2).

**Figure 4.**
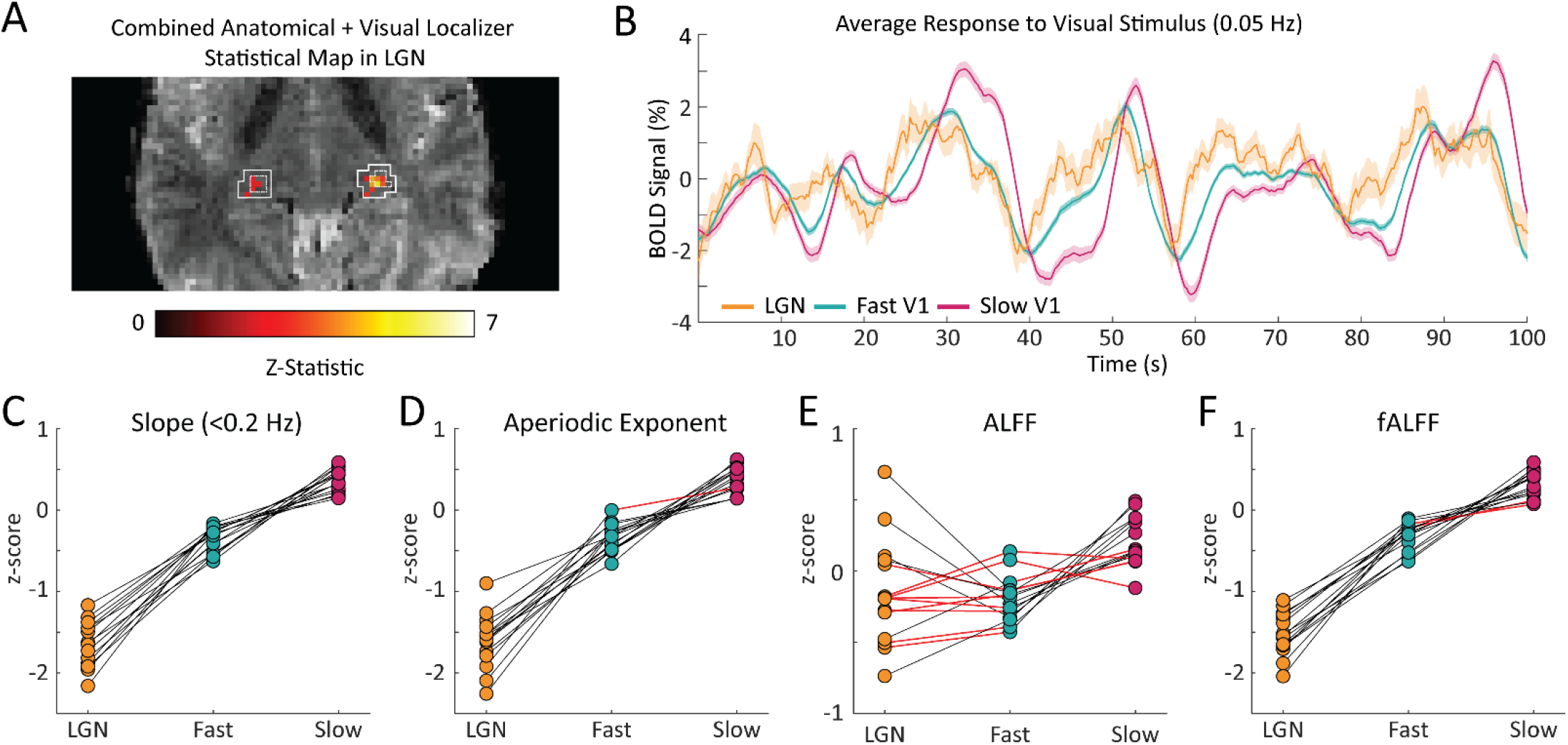
The coupling of response timing and resting-state spectral content is maintained in the LGN. **A)** Example LGN localizer in one subject showing uncorrected z-statistic in the localizer run within the anatomical mask of LGN defined by the white outline. **B)** Average time series of fast and slow V1 voxels compared to LGN voxels for an example subject. LGN displayed a fast visually-driven response, leading even the earliest cortical voxels. Time series are smoothed for display using a 10-point moving average. Shading represents standard error. **C-F)** For each subject we calculated the four resting-state spectral features for the LGN and compared them to the fast and slow voxels in V1. For the **C)** slope, 15/15 subjects showed significant differences between fast vs. LGN and slow vs. LGN; **D)** exponent of an aperiodic 1/f fit, 15/15 subjects showed significant differences between fast-LGN and slow-LGN; **E)** ALFF, 5/15 subjects showed significant differences between fast vs. LGN and 7/15 between slow vs. LGN; and **F)** fALFF, 15/15 subjects showed significant differences between fast-LGN and between slow-LGN. Black lines indicate a significant difference (Wilcoxon rank-sum test, p < 0.05) and red lines indicate a non-significant difference.

### Resting-state spectral information better characterizes neurovascular coupling delays than a breath hold task

Perhaps the most established method of mapping hemodynamic latencies across the brain is using a breath hold task to quantify cerebrovascular reactivity (38, 42–45, 48). Therefore, we tested whether similar information about hemodynamic latencies found using the resting-state spectra could be found in data from the breath hold task. To determine this, we first mapped the vascular latency in response to the breath hold task on a voxel-wise basis across the brain (38). We then compared each voxel’s relative vascular latency from the breath hold task across the three groups identified in the visual task – the fast and slow cortical voxels and LGN voxels. We found that some subjects showed the expected temporal sequence of activation, where LGN voxels and fast cortical voxels respond earlier than slow cortical voxels (Fig. 5A), but this effect was not present in all subjects (Fig. 5B). Even among the subjects that exhibited the expected order of activation, few of them showed individual-level significant differences in breath hold latency between the groups (Fig. 5C). Specifically, in only 7/15 subjects was the average breath hold latency of fast cortical voxels significantly faster compared to slow cortical voxels. Furthermore, the breath hold latency in LGN voxels was slower than expected in some subjects: it was significantly slower than fast cortical voxels in 5 subjects and significantly slower than slow cortical voxels in 1 subject (Wilcoxon rank-sum test, p<0.05). These results demonstrate that although the vascular latencies derived from the breath hold task do show significant differences between fast and slow cortical voxels and LGN voxels across the group, this effect is less robust in individual subjects than the resting-state spectral features. Additionally, not all subjects demonstrated the expected order of latencies between the three groups – with LGN first followed by fast cortical and, lastly, slow cortical voxels. Taken together, these results suggest that the features of the resting-state spectrum capture additional information about local differences in neurovascular coupling delays.

**Figure 5.**
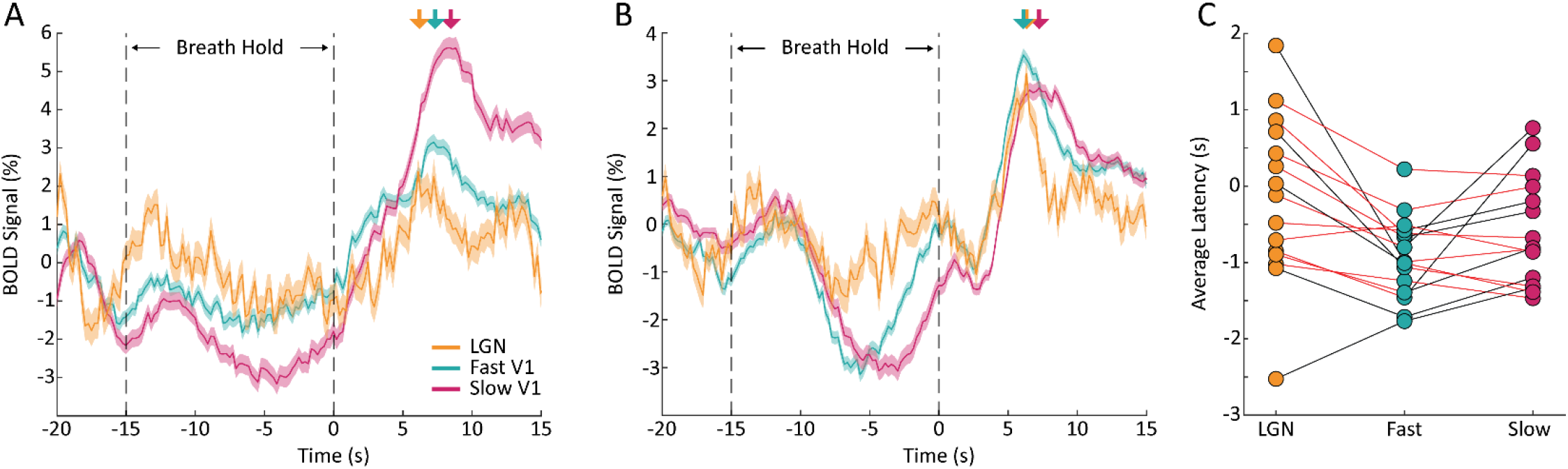
Breath hold vascular latencies yield less robust characterization of task-driven hemodynamic response lags. **A)** Plot of one subject’s average BOLD response to the breath hold task in fast (teal) and slow (pink) cortical voxels as well as LGN voxels (yellow). The shaded areas are the standard error across voxels of that group. Colored arrows denote the peak of the response to the breath hold. The LGN time series peaks slightly earlier than the fast cortical voxels and both the LGN and fast cortical voxels peak well before the slow cortical voxels. This sequence of activation matches what is expected based on the hemodynamic lags across these structures. **B)** Plot of one subject’s average BOLD response to the breath hold task where the order of activation is not as expected. While LGN peaked earliest as expected, the average response in slow cortical voxels peaked before the average response in fast cortical voxels, meaning that the breath hold would not accurately predict latency in this subject. **C)** Comparison of the average breath hold latency in fast, slow, and LGN voxels in all subjects. For 9/15 subjects, the average latency of the fast cortical voxels was less than the slow cortical voxels, as expected, and 7 of these 9 had a significant difference. For only 2/15 subjects, the average latency of the LGN voxels was less than fast voxels, as expected, but only 1 of these 2 had a statistically significant difference. For 4/15 subjects, the average latency of slow cortical voxels was larger than LGN voxels, and 1 of these 4 had a statistically significant difference. (Wilcoxon rank-sum test, p < 0.05).

### Features of the resting-state spectrum can predict voxels with fast or slow hemodynamic response timing

Our results showed consistent signatures of hemodynamic response latency in the resting-state fMRI signal, suggesting that this information could potentially be used to predict local neurovascular latencies. Indeed, we found that these spectral features were significantly correlated with the absolute timing of their hemodynamic responses (Supplementary Fig. S3, Supplementary Table S3). To investigate the utility of the resting-state spectrum to predict the temporal dynamics of the HRF, we tested whether support vector machines (SVMs) could classify slow, fast, and fastest (LGN) voxels using information from the resting-state spectrum. First, we trained a SVM to classify slow, fast, and LGN voxels based on the four features of the resting-state spectra identified in Fig. 3. We found that the classifier validation accuracies, both within each individual subject and on the dataset that combined all subjects, were well above chance (Fig. 6), demonstrating robust prediction of local hemodynamic delays. We next investigated whether our chosen features were sufficient for classification, or whether additional useful information was present in the full resting-state spectrum, by training a second SVM classifier to use each voxel’s resting-state spectrum as the predictor for the classifier. The input to the classifier was the resting-state spectrum limited to up to 0.5 Hz to reduce the number of features fed into the model to avoid overfitting. Once again, we found that the accuracies, both within individual subjects and on the dataset that combined all subjects, were well above chance (Fig. 6; per-subject accuracy in Table S2). The SVM classifiers trained using the spectrum trended toward slightly higher classification accuracies than the pre-selected features (Fig. 6), but the performance of the two models was not significantly different (p=0.2828; Wilcoxon sign-rank test). Regression models aiming to predict specific response timing (rather than classifying fast vs. slow) also performed significantly above chance, but were less reliable than classifying voxels by relative timing (coefficient of determination=0.17; RMSE=0.949; Supplementary Table S4), likely due to the fact that absolute HRF timing depends on task state (27). We therefore concluded that our constructed spectral features are capturing key aspects of the resting-state spectrum that are important for predicting hemodynamic latency.

**Figure 6.**
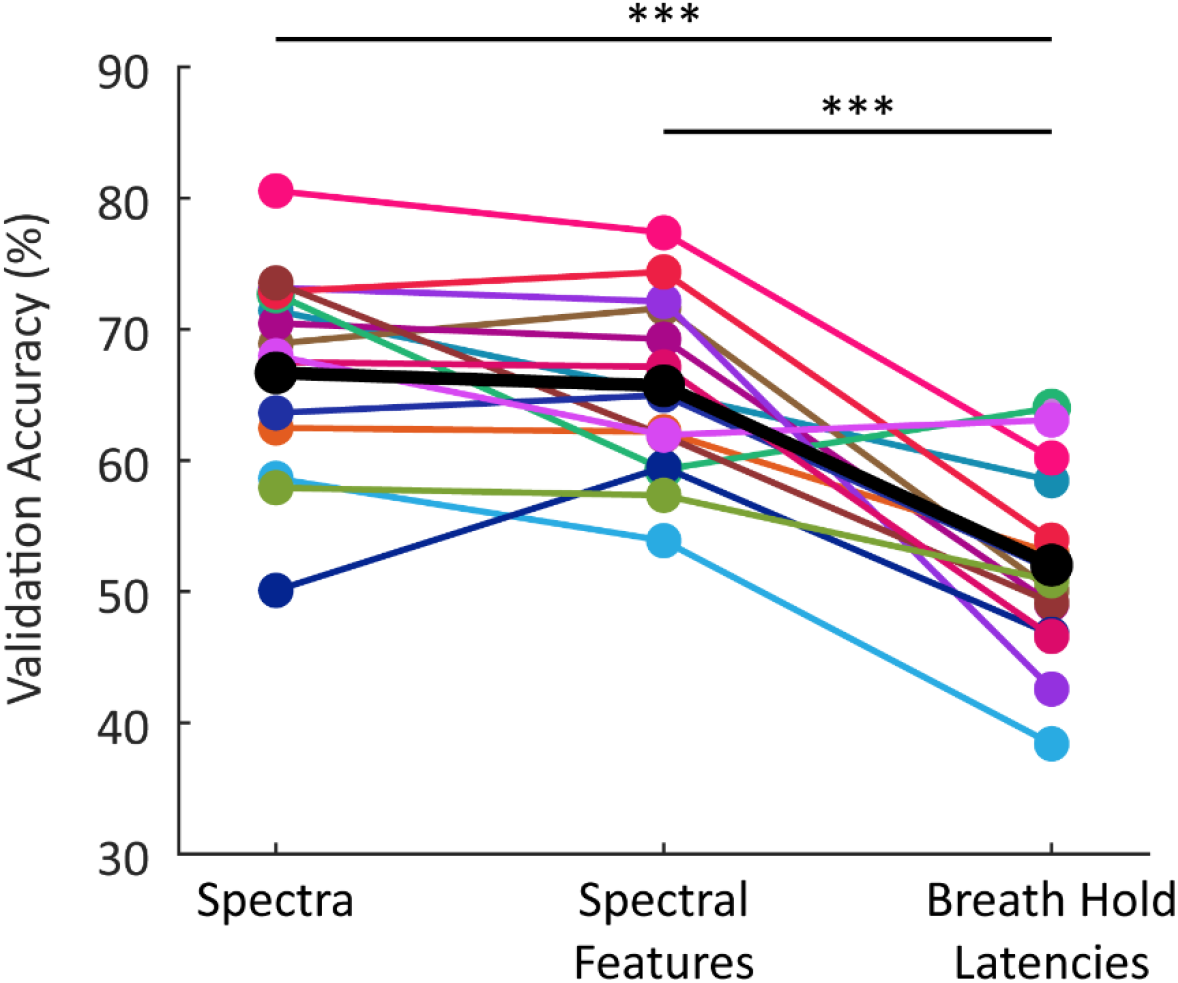
Average classification accuracy of SVM classifier trained using different features. The classification accuracies reported are the average validation accuracy across 1000 bootstraps. The black markers indicate the models trained using resting-state spectral features from all subjects combined into one model; the colored markers show results from individual subjects (n=15). Both models trained using features of the resting-state spectrum performed significantly better than the model trained using the breath hold latencies; however, there was no significant difference in performance between the two models trained using information from the resting-state spectrum (p=0.283). All classifiers perform significantly above chance (33%). *** p<0.0005 (Wilcoxon signed-rank test).

We next tested whether the resting-state signals performed better than the breath hold task at differentiating between fast, slow, and LGN voxels. We trained another SVM classifier to use the breath hold latencies as predictors and found that this model performed significantly above chance (Fig. 6), consistent with prior work demonstrating its utility in predicting delays. However, it nevertheless performed significantly worse than both resting-state models (Wilcoxon sign-rank test, p<0.0005; per-subject accuracy in Supplementary Table S2). This result further supported the conclusion that resting-state spectral information performs well at capturing differences in hemodynamic response timings.

## Discussion

We conclude that the frequency spectrum of resting-state fMRI signals contains rich information about local hemodynamic timing. Our simulations first illustrated the rationale for this effect: HRFs with faster temporal dynamics produce less power in the low frequency bands and a shallower attenuation of power at high frequencies. This effect is preserved even when we account for the higher amplitude of slower HRFs and the 1/f decay of the amplitude of spontaneous neural activity. Following this observation, we constructed several quantitative features based on the resting-state spectrum, each capturing a similar property: the relative amplitude of high vs. low frequencies. These spectral features showed significant differences between fast and slow voxels, and when used as predictors for a SVM were able to classify individual voxels as fast or slow. Our findings demonstrate that the temporal properties of the HRF affect the spectral features of resting-state fMRI signals and present a framework for characterizing the temporal properties of the hemodynamic response across voxels, which is crucial for accurate fMRI analyses.

The work presented here has broad implications for fMRI studies using the frequency content of the BOLD signal to make inferences about intrinsic brain activity. Prior work has identified changes in the spectral content of fMRI signals in certain clinical populations, including major depressive disorder (65), mild cognitive impairment (66), and Alzheimer’s disease (67), and interpreted these differences as changes in intrinsic brain activity. However, as seen here, differences in hemodynamic response timings will also alter the frequency content of fMRI signals. Furthermore, prior work has shown that intrinsic timescales of BOLD activity vary across brain regions and can be used to predict individual subject patterns (68). Our work suggests hemodynamic differences may contribute to these observations. Therefore, changes in the spectral content of fMRI signals can arise not just from differences in intrinsic brain activity, but could also be indicators of different hemodynamic response timing.

A common theme across the spectral features we constructed was that they were sensitive to the relative contributions of low- and high-frequency power. Two features we selected (the slope of the spectrum and the exponent of the aperiodic 1/f fit) are direct measures of the attenuation in power towards higher frequencies. Similar information is contained in the commonly used metrics ALFF and fALFF (60): ALFF is a marker of the power in low frequency bands and fALFF is a measure of the ratio of high to low frequency power. While each of these features showed significant differences between fast and slow cortical voxels, only the slope and fALFF exhibited significant differences within each individual subject. These features explicitly capture the relative difference in high-frequency vs. low-frequency power, corresponding to the main prediction of our simulations. Conversely, ALFF had the poorest sensitivity within subjects which could be due to the fact that ALFF is the only feature that does not include information from both low and high frequency ranges and is instead limited to a relatively narrow band of frequencies, 0.01-0.08 Hz. Overall, our results suggest that while some information is contained in the magnitude of the fMRI signal alone, capturing the relative power in high versus low frequencies is the most important metric for predicting hemodynamic timing. Moreover, using the resting-state spectrum itself did not perform significantly better than our chosen features in classifying fast and slow voxels, suggesting that the relative power is indeed the primary contributor when predicting variations in local HRF timings.

A unique advantage of fMRI is its ability to image throughout the whole brain, and our results suggest the potential for extracting more information from fMRI studies of subcortex. Due to significant improvements in both the sensitivity and spatial resolution of fMRI, an increasing number of studies are utilizing fMRI to study small, deep brain structures such as the thalamus and brainstem (69–72). The ability to image these deeper brain structures in humans opens the door to studying diverse aspects of cognition associated with these deeper brain regions (69–73). However, these deep brain structures also have unique physiological and anatomical properties that alter their vascular dynamics (61–63, 74, 75). Faster hemodynamic responses are frequently reported in subcortex, but hemodynamic responses in these regions are less well characterized than in the cortex. Our analyses found that variations in HRF timing are reflected in the frequency spectra of thalamic voxels as well, despite their differing signal-to-noise ratio. This result demonstrates the utility of our approach in subcortex and could benefit neuroimaging studies of structures such as the thalamus, a target of increasing interest in fMRI.

Improvements in the temporal resolution of BOLD fMRI have also sparked interest in detecting neural sequences at sub-second timescales, which are highly relevant for many studies of cognition. Recent studies have leveraged fast fMRI to detect rapid sequences of neural events related to visual sequence detection (76), auditory dynamics (77), and changes in arousal state (22). As we continue to identify these rapid neural sequences, it will become even more crucial to consider how the hemodynamic response varies across regions to determine whether a given sequence represents regional differences in neuronal or in hemodynamic timing. Considering spectral signatures can support inference of precise timing of neural activity by providing information about relative hemodynamic latencies between voxels and regions.

We found that the resting-state signals predicted voxel-wise differences in relative hemodynamic response timing significantly better using a breath hold to estimate vascular delays, the current gold standard. This observation could be explained by multiple factors. First, the models trained using the breath hold latencies were only able to leverage one piece of information: the measure of vascular latency derived from cross-correlation. The information from taking the cross-correlation in the time domain may also be more sensitive to noise, especially at the fast sampling rate of our scans. However, a more important factor may be the biological mechanisms generating each signal. The breath hold task directly modulates cerebrovascular reactivity with minimal accompanying changes in CMRO_2_, allowing for assessment of local cerebral vascular reactivity uncoupled from neuronal activation (42). CVR is important to assess in many clinical applications, and the breath hold-based approach remains a gold standard for CVR mapping (42–44, 48). However, while CVR is a significant modulator of the hemodynamic response, it is only one component of neurovascular coupling, and there are many other factors and signaling pathways that also contribute. In particular, local metabolic factors and feedback mechanisms also modulate of blood flow in the brain and are not replicated in the breath hold task (50, 78). The hemodynamic responses induced by neural activity may, therefore, not be identical in timing to those induced by a purely vascular signal. By contrast, the signals obtained from resting-state fMRI are coupled to underlying neural activity in an analogous manner as during a task condition (54, 55, 57). The improved performance of the resting-state-based prediction may therefore reflect that it is intrinsically more similar to a task than the breath hold condition, as the underlying biological origins of resting-state signals share common mechanisms with task-induced neurovascular coupling. Future applications may therefore benefit from continuing to use breath tasks as a gold standard to assess CVR, whereas resting-state analyses may be a better metric of neurovascular coupling.

The work presented here suggests a wide range of neuroscience applications for our approach to measuring hemodynamic timing. One logical next step would be to use the ability to characterize temporal variation in the HRF to not just predict, but to correct for vascular delays. Previous work demonstrated that correcting for varying hemodynamic latencies across the brain can affect functional connectivity analyses (35, 38). Our results could further enhance removal of non-neural latency differences that confound functional connectivity metrics, both static and dynamic, and can increase confidence that the networks we are analyzing are derived from neuronal dynamics. This has become increasingly of interest as more studies use functional connectivity, particularly in resting-state, to study dynamics underlying diseases such as PTSD (79–82), Alzheimer’s Disease (83–86), Parkinson’s Disease (87–91), and others (92). Since changes in neurovascular function have been observed in many disorders (93–95), analyzing spectral dynamics may help interpret functional connectivity differences in clinical populations.

Although we focused on the utility of resting-state spectral information to classify voxels as having relatively fast or slow hemodynamic response timings, we did also examine the correlation with absolute timing measures. We found that most subjects had significant positive correlations between each of the spectral features and the task response latency. Despite this fact, when estimating continuous predictions of absolute hemodynamic response lag, prediction performance was poor. Importantly, previous evidence has shown that the absolute timing of the HRF varies between task and resting-state conditions (27), suggesting that caution is needed in using absolute measures of hemodynamic latencies, as these may not generalize across conditions. Measuring relative differences in HRF timing might therefore be a more effective method to correct for variations in the hemodynamic response across brain regions, since task state can modulate the absolute timing of the hemodynamic response. Furthermore, the relative differences provide the key information necessary to interpret sequences of activity across brain regions, enabling examination of whether purely vascular differences are present across those regions.

Together, our results demonstrate that the resting-state fMRI signal contains information about local hemodynamic response speeds. This approach can help understand brain-wide variations in HRF dynamics, which is critical as the field moves towards a new era of fMRI studies utilizing fast fMRI to study rapid neuronal dynamics and higher-level cognition.

## Acknowledgments

This work was supported by National Institutes of Health grants R00-MH111748, U19-NS123717, and R01-AG070135, the Searle Scholars Program, the Pew Biomedical Scholars Program, the Sloan Fellowship, and the One Mind Rising Star award. Resources were provided by NIH grant P41-EB030006 and T32-GM008764.

## Data sharing statement

Upon acceptance, the data used in this paper will be deposited publicly and freely shared.

## Materials and Methods

### Simulations

Spectra of simulated BOLD responses were generated by convolving a given HRF with oscillating stimuli, ranging from 0.1-0.5 Hz, and taking the magnitude of the simulated BOLD response as the power at that frequency (Fig. 1A). We used six HRFs with varying time-to-peak (TTP), full width at half maximum (FWHM), and peak percent signal changes (PSCs) to represent a range of physiologically relevant HRFs (Fig. 1B). These properties were drawn from previous work characterizing varying HRF temporal dynamics at different cortical depths (11) and the values used are reported below in Supplementary Table S5. We also normalized these HRFs by their maximum percent signal change and re-simulated the BOLD responses to create a new simulated spectrum for each HRF (Fig. 1C). Additionally, we performed simulations to account for the dominant 1/f-like spectral pattern of neural activity by setting the amplitude of the oscillating stimuli to be 1/stimulus frequency ranging from 0.1-0.5 Hz (Fig. S1).

### Subject Population

All experimental procedures were approved by the Massachusetts General Hospital Institutional Review Board and all subjects provided informed consent. 21 participants were scanned in total; 5 were excluded for excessive motion and 1 was excluded for poor performance on the visual task suggesting they had closed their eyes. This left 15 subjects whose data was analyzed (mean age = 28 years, range = 22-42 years, 8 female).

### Experimental Design

Subjects underwent a total of 7 functional scans: 3 visual stimulus, 2 breath hold, and 2 resting-state runs. All stimuli were programmed in MATLAB using Psychtoolbox (96).

### Visual Stimulus

Each visual stimulus functional run lasted 254 seconds, with the first 14 seconds showing a gray screen with the red fixation dot and the following 240 seconds consisting of the 12-Hz counterphase flickering radial checkerboard. To drive continuous neural oscillations in the visual cortex, the luminance contrast of the flickering checkerboard oscillated at a frequency of 0.05 Hz (except for one subject who was presented with 0.1 Hz oscillations). To assist the subjects with fixation, in the center of the visual field was a red dot that changed brightness at random intervals. Subjects were directed to press a button whenever the brightness of the red dot changed, and their average response time and response accuracy was reported at the end of each run. This allowed us to monitor participant engagement with the task. Each subject participated in 3 visual stimulus runs.

### Breath Hold Task

For each breath hold run subjects performed 8 repetitions of an adapted version of a previously established breath hold task (38): a block comprised of 27 seconds of free breathing, 3 cycles of paced breathing (3 seconds breathe out, 3 seconds breathe in), a 15 second breath hold, and, lastly, a 30 second period of free breathing. The total time for the breath hold task scans was 8.5 minutes. The instructions for breathing were projected to the subjects displaying the text “Breathe Freely”; “Breathe Out”; “Breathe In”; and “HOLD BREATH.” All but one subject participated in 2 breath hold runs; a single subject performed only 1 breath hold run. For 2 subjects, a single breath hold run was excluded from analyses for excessive motion defined here as greater than 0.5 mm average motion across the whole run. No additional runs were excluded for these subjects.

### Resting-state

Each subject participated in 2 resting-state runs where they were instructed to relax in the scanner with their eyes open and to try not to fall asleep. For some of the subjects, a fixation dot was presented to help minimize eye movements. Each resting-state scan lasted 8.5 minutes.

### MRI Data Acquisition

Subjects were scanned on a 7-Tesla Siemens MAGNETOM scanner with a custom-built 32-channel head coil. Anatomical images were acquired with 0.75-mm isotropic multiecho magnetization-prepared rapid gradient-echo (MEMPRAGE) protocol (97) with TR = 2,530 ms, echo time (TE) = 1.76 ms and 3.7 ms, inversion time (TI) = 1100 ms, echo-spacing = 6.2 ms, 7° flip angle, bandwidth = 651 Hz, in-plane acceleration R=2, FOV = 320 x 320 x 244 mm and a total scan time of 7:20 minutes. For functional runs, 15 oblique slices were positioned to target the calcarine sulcus to include primary visual cortex (V1) and angled to include the lateral geniculate nucleus (LGN) located in the thalamus. Functional runs were acquired as single-shot gradient-echo EPI with 2 mm isotropic resolution, TR = 227 ms, TE = 24 ms, echo-spacing = 0.59 ms, 30° flip angle, bandwidth = 2604 Hz, in-plane acceleration R = 2, SMS Multiband Factor = 3, CAIPI shift=FOV/3 (98).

### Physiological Monitoring

For all the functional scans, subjects’ heart rate and respiration were monitored using piezoelectric transducer on the non-dominant thumb and a respiratory belt around the upper rib cage, respectively. The physiological recordings were obtained at a sampling rate of 100 Hz using a PowerLab physio box connected to a computer running LabChart 7 from ADInstruments.

### fMRI Analyses

#### fMRI Preprocessing

Anatomical images were bias-corrected using SPM (https://www.fil.ion.ucl.ac.uk/spm/) and segmented using FreeSurfer (64). Functional runs were preprocessed with slice-timing correction, performed using FSL (http://fsl.fmrib.ox.ac.uk/fsl/fsl-wiki/), and motion correction, performed using AFNI software (https://afni.nimh.nih.gov/). No spatial smoothing was applied.

Because fast fMRI has distinct contributions from systemic physiological noise, including cardiac rhythms and respiration, physiological noise removal was performed on the visual stimulus and resting-state functional runs using dynamic regression adapted from RETROICOR (99) in runs where physiological recordings were successfully collected. In runs where physiological recordings were not successfully collected (2 runs total), physiological noise was removed using a statistical model of harmonic regression with autoregressive noise (HRAN) (100).

#### Visual Localizer

For each subject, one of the visual stimulus runs was used as a functional localizer to identify voxels that were significantly driven by the oscillating stimulus. A general linear model (GLM) was fit in FSL (http://fsl.fmrib.ox.ac.uk/fsl/fslwiki/) using sine and cosine basis functions with the same period as the stimulus. The F-statistic of the combined fit to both the sine and cosine basis function was transformed to a Z-score and voxels with a Z-score above 2.5 were selected for further analysis. This functional localizer was then constrained by its intersection with the anatomical definition of V1 (Fig. 2B). Specifically, the V1 segmentation was generated automatically from the MEMPRAGE volume based on the cortical surface reconstruction generated using FreeSurfer (64). The selected voxels from the localizer run were then mapped to each other functional run in a single transformation step. This was done by first registering all functional runs to the anatomical scan using boundary-based registration (101) and then resampling the desired volume into the localizer field of view using the registration matrices.

#### LGN Segmentation

The lateral geniculate nucleus (LGN) was segmented using both anatomical and functional constraints. The anatomically defined boundaries of the LGN were generated using the FreeSurfer developmental version that generates an individual-level probabilistic atlas in individual anatomical space (102). From this probabilistic atlas we considered voxels with at least a 30% probability of falling within LGN and dilated this mask to capture border voxels. We next applied a functional constraint using the visual localizer where voxels in the dilated mask with a Z-score above 2.5 were considered part of our final LGN map in each subject.

#### Voxel-wise Phase Analysis and Groupings

We then averaged the two remaining visual stimulus runs and extracted voxel’s average time series between the two runs. We discarded the first 14 seconds to analyze the steady-state response to the visual stimulus. An estimation of the voxel’s lag in relation to the oscillating stimulus was calculated using the arctangent of the sine and cosine regressor estimates. This allowed us to generate a histogram of latencies to the visual stimulus (Fig. 2C). To extract “fast” and “slow” reacting voxels, a Gaussian model was fit to the histogram of phase delays. The centroid (*b*) and Full Width at Half Maximum (*FWHM*) of the Gaussian fit were calculated. Groups of fast and slow voxels were then Edges of the Gaussian fit were defined as 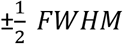 and fast and slow groups were made that each had a width of 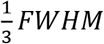. Fast voxels were identified as being within 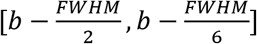 while slow voxels were identified as 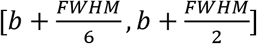 (Fig. 2C). This procedure was done for each individual subject and on average yielded 142 fast voxels (range 64-236) and 139 slow voxels (range 68-254). Masks of fast and slow voxels were generated per subject and then mapped to the resting-state runs to inform the spectral analysis (Fig. 2D).

#### Resting-state Spectral Analysis

Fast and slow voxels were always identified in task-driven runs, allowing us to assess frequency content in the resting-state run using fully independent data. The maps of fast and slow voxels were registered to each individual resting-state run, and for each voxel within these masks, after discarding the first 14 seconds, the voxel-wise resting-state power spectrum was calculated using the Chronux toolbox (103) with five tapers. We used 4 features to characterize the resting-state spectra: (1) slope of linear fit under 0.2 Hz; (2) the exponent of the aperiodic 1/f fit under 0.5 Hz; (3) the amplitude of low frequency fluctuations (ALFF); and (4) the fractional ALFF (fALFF). Each of these features was z-scored within the run and then averaged on a voxel-wise basis between the two resting-state runs. All analysis of the resting-state spectra was performed in MATLAB. See Fig. 3A-D for more information.

##### Slope

A linear fit was generated for each voxel’s resting-state spectrum under 0.2 Hz using least-squares to determine the coefficients of a first order polynomial. From this we were able to record the slope of that linear fit for each voxel.

##### Exponent of Aperiodic Fit

Equation 1 was fit using the Levenberg-Marquardt algorithm to solve non-linear least squares for each voxel’s resting-state spectra. In this equation, *F* is the independent variable and the *b* and *x* are the values being fit. The exponent of the resultant fit (*x*) was recorded for each voxel.

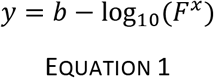

##### Amplitude of Low Frequency Fluctuations (ALFF)

For each voxel, ALFF was calculated according to the method outlined in (60). Briefly, each voxel’s time series was band pass filtered between 0.01 – 0.08 Hz. Then, the voxel’s time series is transformed into the frequency domain via Fast Fourier Transform (FFT). The power at each frequency is proportional to the square of the amplitude of the FFT at that frequency, and for each voxel, the ALFF value was taken as the averaged square root of the power in the 0.01-0.08 Hz frequency range. This is shown in Equation 2 where *FFT*(*k*) is the magnitude of the FFT at frequency *k*, *N* is the number of time points, *P* is the power spectrum, and the bar represents the average in the specified range.

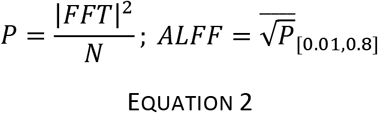

##### Fractional ALFF (fALFF)

Each voxel’s fALFF was calculated as described in (60). Fractional ALFF is briefly defined as the ratio of the power of each frequency at the low frequency range (0.01-0.08 Hz) to that of the “global” frequency range (0.01-0.25 Hz). See Equation 3.

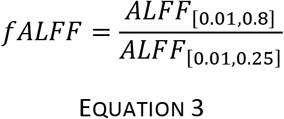

#### Breath-Hold Latency Calculations

Voxel-wise hemodynamic latencies were calculated according to the method outlined in (38). Each voxel’s hemodynamic latency was defined as the time-lag yielding the maximum cross-correlation between the given voxel’s time series, *x*(*t*), and a reference time series, *y*(*t*). The reference time series was found by taking the average time series across voxels in the brain that exceeded a minimum correlation of *r* > 0.25 with the breath hold task regressor. This breath hold task regressor was defined as the convolution of a box car function, where the value is set to 1 during the breath hold and 0 at other times, and a sign-reversed canonical HRF (31). Both *x*(*t*) and *y*(*t*) were resampled to a resolution of 100 ms before computing cross-correlations.

#### Support Vector Machine (SVM) Classification of Fast, Slow, and LGN Voxels

SVM classifiers were trained both within and across subjects using Scikit-learn in Python (104). Three models were trained with different predictive features: (1) the resting-state spectrum between 0 to 0.5 Hz, (2) the 4 features of the resting-state spectrum previously identified, and (3) the latency of the response to the breath hold task. The resting-state spectrum was extracted by taking the power at frequencies up to 0.5 Hz ultimately generating a set of 461 features. For all models, before being put into the classifier, the data was normalized by removing the mean and dividing by the standard deviation across voxels for each feature independently using the StandardScaler function of Scikit-learn. For all models the parameters of the SVM classifier were as follows: regularization parameter (*C*) = 10, kernel type = radial basis function (rbf), kernel coefficient 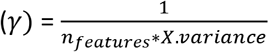. To get validation accuracies both within and across subjects, 1000 bootstraps were performed where the 80-20 test-train split of voxels was randomly chosen for each bootstrap. The average validation accuracies over the 1000 bootstraps were calculated along with the 95% confidence intervals. This methodology was followed for all 3 SVM classifiers.

#### Support Vector Machine (SVM) Regression to Predict Relative Hemodynamic Response Latency

SVM models for regression were trained within subject to continuously predict the hemodynamic response latency estimated from the visual stimulus. We limited the regression to voxels whose hemodynamic response latency relative to the median was between [-3, 3] sec. Two different models were trained with different predictive features: (1) the resting-state spectrum between 0 to 0.5 Hz and (2) the 4 features of the resting-state spectrum previously identified. The input data to the model was normalized by removing the mean and dividing by the standard deviation cross voxels using the StandardScaler function of Scikit-learn. The parameters of the SVM regression model were as follows: kernel type = radial basis function, kernel coefficient 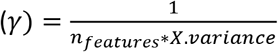, regularization parameter = 10, epsilon = 0.1. To get the validation coefficient of determination (*R*^2^), 1000 bootstraps were performed on an 80-20 test-train split of voxels that were randomly chosen for each bootstrap. The average *R*^2^ over the 1000 bootstraps was calculated along with the 95% confidence interval.

## Supplementary Material

**Supplementary Figure S1.**
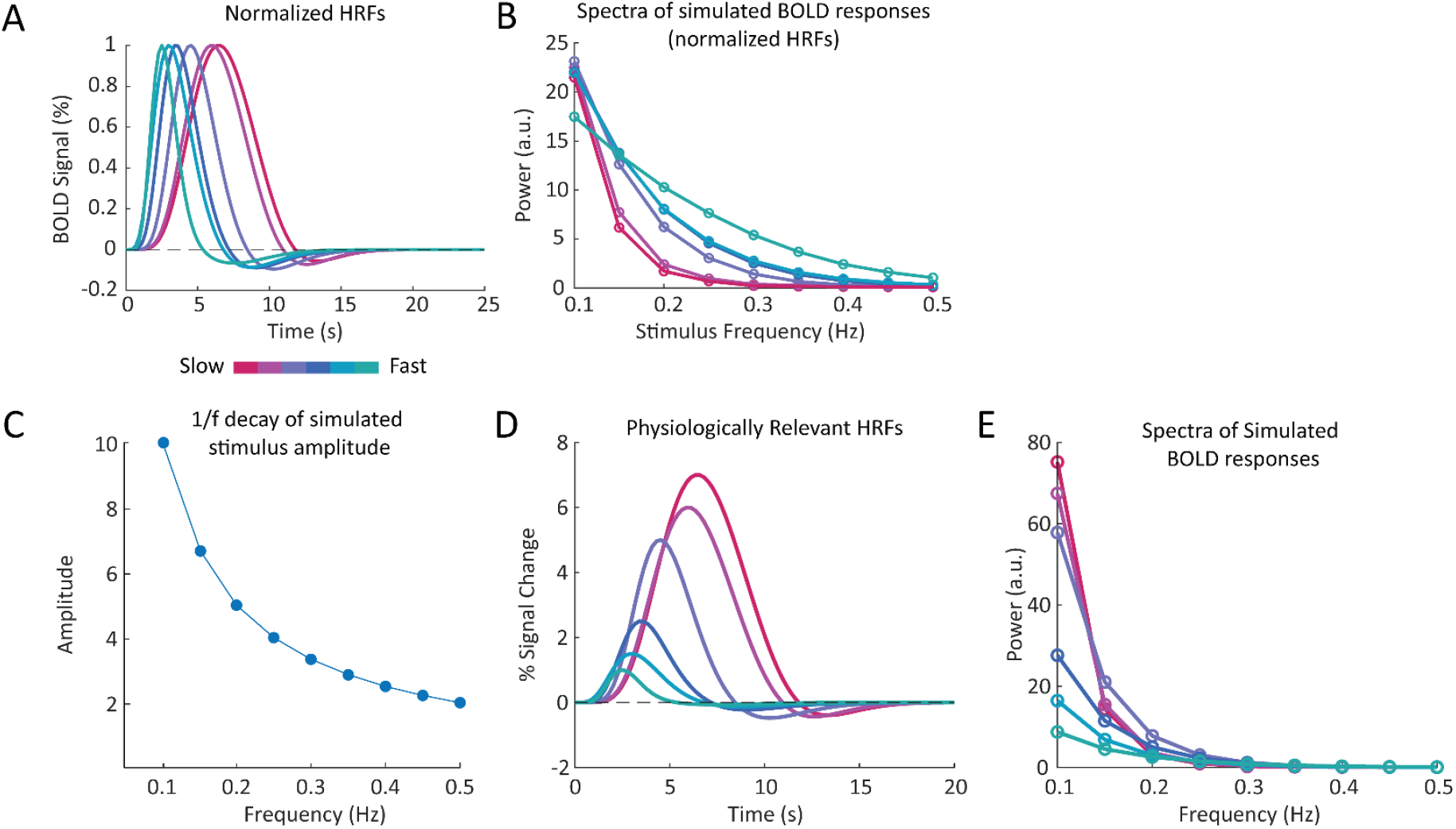
Simulation results are robust to changes in HRF amplitude and 1/f decay of stimulus amplitude. **A)**We normalized each HRF to its respective peak amplitude to examine effect on frequency spectra. B)The differences in spectral content of the BOLD signal are not simply due to the amplitude of each HRF but rather a signature of the temporal dynamics of the HRF, as the slope differences persist when normalizing for amplitude, with slower HRFs generating steeper frequency responses. **C)**Plot showing the magnitudes of the neural waveform at each frequency and their 1/f decay. **D)**Physiologically relevant HRFs used in simulations. **E)** Temporal properties of HRF noticeably affect the slope of the simulated spectra even when oscillation amplitudes decay with 1/f pattern, demonstrating that distinct slopes are expected for distinct HRFs regardless of the specific neural signal amplitude.

**Supplementary Figure S2.**
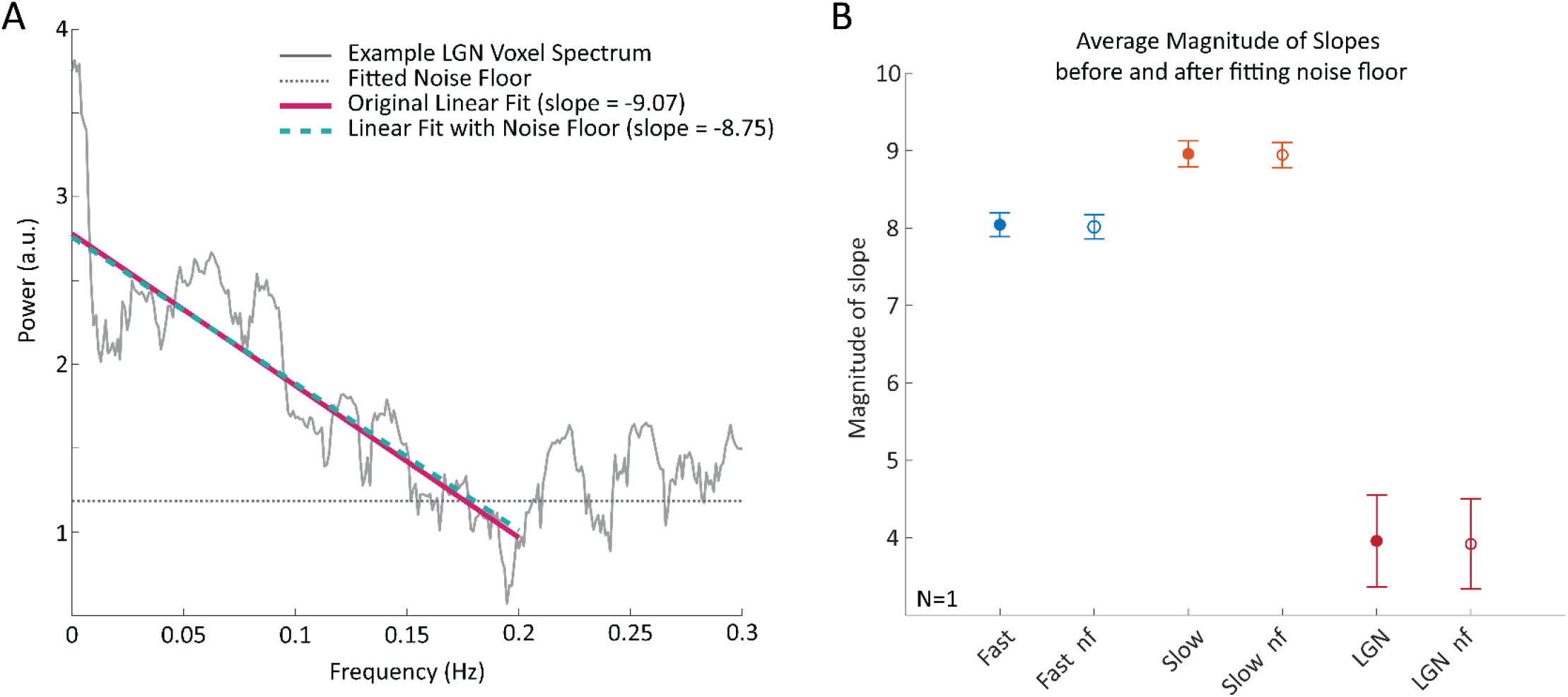
Accounting for thermal noise does not significantly change the estimated slope of the frequency spectrum under 0.2 Hz in V1 or LGN voxels. **A)** Spectra of example LGN voxel demonstrating the method for accounting for the noise floor in the slope of the linear fit. The original linear fit and the linear fit with the noise floor produce similar results. **B)** Average magnitude of slopes before and after fitting the noise floor in fast and slow cortical voxels as well as LGN voxels with error bars showing SEM. The within group averages did not significantly change when fitting for the noise floor versus not.

**Supplementary Figure S3.**
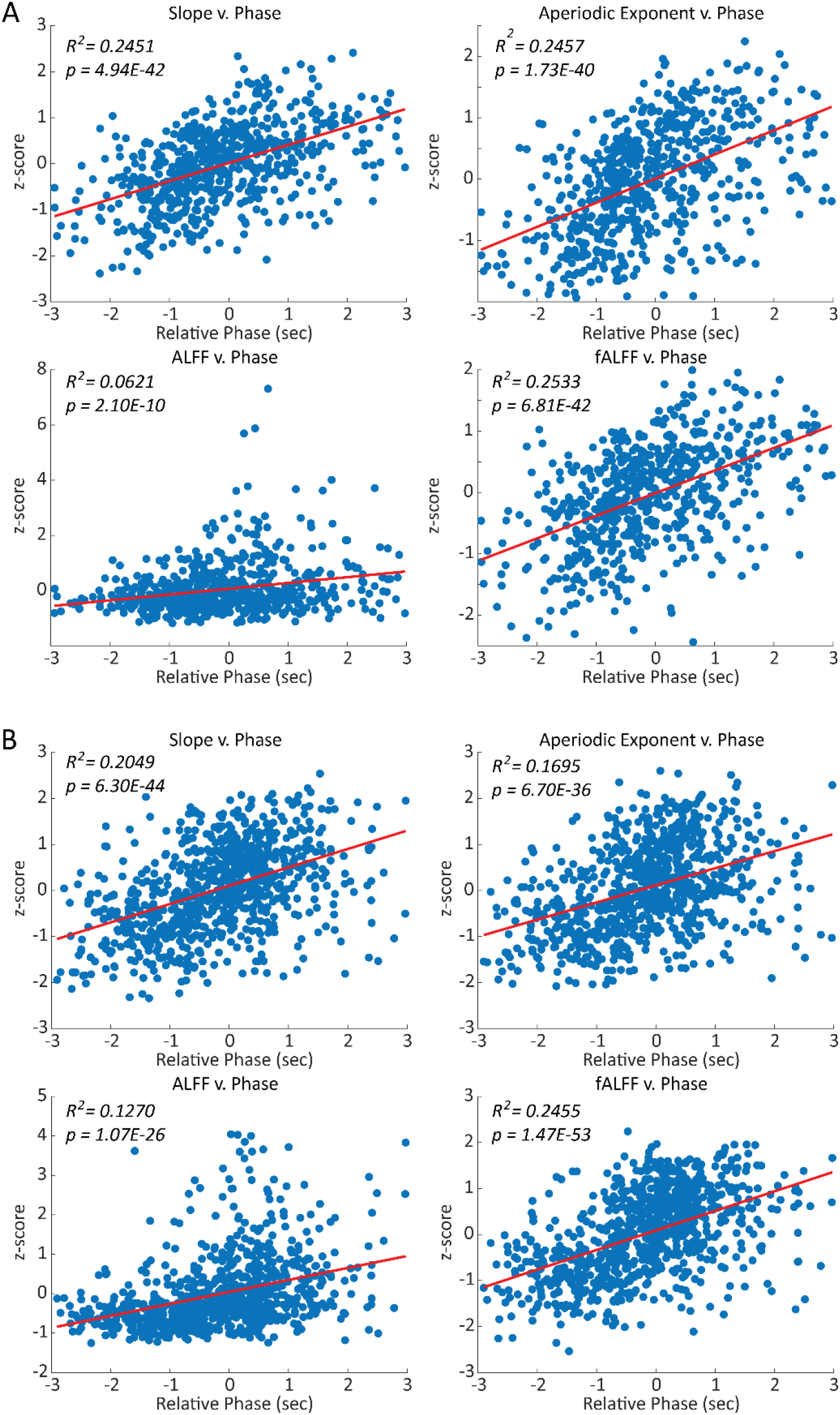
Example subjects showing significant (p<0.05) correlations between each spectral feature and phase on a voxel-wise basis. Each point represents a single voxel, red-line shows linear fit. p-values are from a linear hypothesis test on the model coefficients. **A)** Example subject showing significant, positive correlations between each spectral feature and the magnitude of the phase. **B)** A second example subject also showing a significant, positive correlation between each spectral feature (except ALFF) and the magnitude of the phase.

**Table S1:** p-values for all subjects and all resting-state spectral features between fast, slow, and LGN voxels, within individual subjects. Significant differences based on Wilcoxon rank-sum test (p<0.05) are bolded.

|  | *Slope* < 0.2 Hz |  |  | *Aperiodic Exponent* |  |  | *ALFF* |  |  | *fALFF* |  |  |
| --- | --- | --- | --- | --- | --- | --- | --- | --- | --- | --- | --- | --- |
|  | Fast-Slow | Fast-LGN | Slow-LGN | Fast-Slow | Fast-LGN | Slow-LGN | Fast-Slow | Fast-LGN | Slow-LGN | Fast-Slow | Fast-LGN | Slow-LGN |
| S1 | **1.13E-14** | **6.51E-09** | **7.15E-17** | **1.13E-12** | **3.26E-09** | **1.71E-16** | **6.98E-14** | 0.7237 | **7.52E-09** | **1.64E-14** | **7.88E-10** | **7.38E-17** |
| S2 | **1.29E-21** | **3.93E-10** | **6.16E-14** | **6.21E-22** | **6.12E-11** | **9.59E-14** | **2.33E-10** | **0.0021** | 0.4402 | **2.59E-20** | **3.26E-05** | **1.14E-10** |
| S3 | **5.06E-16** | **3.08E-06** | **1.13E-10** | **7.46E-17** | **4.79E-06** | **5.69E-11** | **4.89E-05** | 0.8869 | **0.0159** | **3.34E-15** | **7.78E-07** | **6.76E-11** |
| S4 | **3.51E-03** | **4.42E-06** | **2.06E-08** | **3.95E-03** | **3.82E-07** | **2.59E-09** | **3.20E-03** | 0.2091 | **0.0079** | 0.0643 | **2.09E-05** | **2.05E-07** |
| S5 | **6.67E-08** | **2.88E-09** | **6.29E-10** | **3.91E-12** | **1.17E-09** | **3.45E-10** | **1.15E-05** | **2.98E-05** | **0.0446** | **2.02E-06** | **6.62E-09** | **8.86E-10** |
| S6 | **3.61E-25** | **1.01E-10** | **1.85E-12** | **4.20E-22** | **1.12E-10** | **1.34E-12** | **3.59E-20** | 0.9969 | **6.20E-06** | **3.72E-30** | **9.55E-07** | **1.27E-11** |
| S7 | **9.71E-04** | **2.51E-04** | **7.79E-05** | 0.1432 | **2.51E-04** | **9.19E-05** | 0.5493 | **0.0498** | 0.0867 | **0.0322** | **3.76E-04** | **1.75E-04** |
| S8 | **6.89E-14** | **1.39E-09** | **4.15E-14** | **1.20E-20** | **1.74E-09** | **1.41E-14** | **3.14E-03** | 0.4995 | 0.2247 | **4.56E-11** | **1.48E-09** | **9.87E-14** |
| S9 | **1.72E-24** | **9.55E-15** | **7.08E-21** | **6.84E-18** | **1.64E-14** | **6.48E-21** | **0.5928** | 0.6579 | 0.2915 | **1.27E-24** | **5.91E-11** | **8.89E-20** |
| S10 | **4.47E-10** | **1.75E-13** | **9.63E-19** | **9.19E-04** | **9.38E-18** | **9.81E-22** | 0.1464 | 0.3322 | 0.8048 | **9.32E-10** | **5.00E-13** | **5.58E-18** |
| S11 | **6.65E-04** | **2.19E-11** | **6.32E-12** | **4.96E-04** | **1.39E-11** | **4.80E-12** | **6.78E-04** | 0.0585 | 0.1637 | 0.0533 | **1.05E-09** | **4.39E-10** |
| S12 | **4.08E-13** | **1.41E-10** | **3.68E-13** | **1.77E-15** | **4.72E-09** | **1.01E-12** | **1.83E-09** | **5.88E-08** | **1.15E-10** | **1.83E-07** | **1.03E-09** | **2.38E-12** |
| S13 | **8.83E-10** | **7.77E-08** | **1.79E-11** | **5.32E-09** | **4.31E-04** | **8.21E-09** | 0.3715 | **4.92E-10** | **5.66E-07** | **8.67E-08** | **9.20E-10** | **2.47E-12** |
| S14 | **5.76E-04** | **4.19E-06** | **1.54E-08** | **2.70E-03** | **5.63E-06** | **8.22E-09** | 0.3232 | 0.1669 | 0.3733 | **0.0233** | **5.27E-06** | **9.18E-08** |
| S15 | **1.92E-12** | **1.06E-07** | **9.38E-11** | **6.07E-14** | **1.44E-08** | **2.79E-11** | **1.81E-07** | **8.08E-04** | 0.8278 | **8.47E-13** | **8.46E-05** | **2.32E-09** |

**Table S2:** Average Classification Accuracies per subject. Results are average of over 1000 bootstraps with 95% confidence intervals for each subject on each model trained. For each subject the model with the highest accuracy is bolded. Chance is 33%.

|  | Subsampled Spectra | Spectral Features | Breath Hold Latencies |
| --- | --- | --- | --- |
| S1 | **71.43 % (0.31)** | 65.21 % (0.37) | 58.46 % (0.36) |
| S2 | 68.91 % (0.31) | **71.63 % (0.31)** | 50.08 % (0.34) |
| S3 | **70.46 % (0.44)** | 69.25 % (0.46) | 49.06 % (0.50) |
| S4 | **58.63 % (0.49)** | 53.88 % (0.47) | 38.39 % (0.45) |
| S5 | **62.48 % (0.30)** | 62.19 % (0.31) | 52.94 % (0.35) |
| S6 | **80.56 % (0.25)** | 77.37 % (0.29) | 60.19 % (0.34) |
| S7 | **72.69 % (0.48)** | 59.31 % (0.54) | 64.01 % (0.53) |
| S8 | **73.18 % (0.35)** | 72.12 % (0.37) | 42.57 % (0.38) |
| S9 | 72.87 % (0.29) | **74.38 % (0.29)** | 53.93 % (0.34) |
| S10 | **73.54 % (0.24)** | 61.90 % (0.26) | 49.16 % (0.26) |
| S11 | 50.09 % (0.44) | **59.49 % (0.43)** | 46.74 % (0.45) |
| S12 | 63.63 % (0.33) | **64.98 % (0.34)** | 51.65 % (0.37) |
| S13 | **67.50 % (0.32)** | 67.14 % (0.35) | 46.58 % (0.35) |
| S14 | **57.92 % (0.37)** | 57.34 % (0.38) | 50.83 % (0.36) |
| S15 | **67.99 % (0.35)** | 61.94 % (0.36) | 63.08 % (0.39) |
| COMBINED | **66.68 % (0.08)** | 65.70 % (0.08) | 52.02 % (0.11) |

**Table S3.** Subject-wise results from fit of linear model relating each spectral feature with phase. p-values are from a linear hypothesis test on the model coefficients.

|  | Slope < 0.2 Hz |  |  | Aperiodic Exponent |  |  | ALFF |  |  | *f*ALFF |  |  |
| --- | --- | --- | --- | --- | --- | --- | --- | --- | --- | --- | --- | --- |
|  | x1 | R<sup>2</sup> | p-value | x1 | R<sup>2</sup> | p-value | x1 | R<sup>2</sup> | p-value | x1 | R<sup>2</sup> | p-value |
| S1 | 0.3865 | 0.1058 | 1.51E-16 | 0.3391 | 0.0781 | 1.93E-12 | 0.3998 | 0.1063 | 1.28E-16 | 0.3868 | 0.1080 | 7.14E-17 |
| S2 | 0.3657 | 0.1204 | 1.46E-20 | 0.4044 | 0.1251 | 2.40E-21 | 0.2475 | 0.0500 | 4.24E-09 | 0.3239 | 0.1017 | 1.93E-17 |
| S3 | 0.3935 | 0.2451 | 4.94E-42 | 0.3940 | 0.2457 | 1.73E-40 | 0.2096 | 0.0621 | 2.10E-10 | 0.3691 | 0.2533 | 6.81E-42 |
| S4 | 0.2592 | 0.1153 | 5.87E-17 | 0.2692 | 0.1208 | 9.75E-18 | 0.1670 | 0.0351 | 6.14E-06 | 0.2188 | 0.0922 | 1.04E-13 |
| S5 | 0.1836 | 0.0413 | 1.93E-09 | 0.2591 | 0.0718 | 1.49E-15 | 0.1394 | 0.0227 | 9.65E-06 | 0.1727 | 0.0364 | 1.81E-08 |
| S6 | 0.3990 | 0.2049 | 6.30E-44 | 0.3721 | 0.1695 | 6.70E-36 | 0.3041 | 0.1270 | 1.07E-26 | 0.4249 | 0.2455 | 1.47E-53 |
| S7 | 0.2918 | 0.1363 | 1.09E-27 | 0.2558 | 0.0911 | 1.34E-18 | 0.1839 | 0.0710 | 1.09E-14 | 0.3188 | 0.1625 | 3.73E-33 |
| S8 | 0.2714 | 0.1870 | 8.47E-37 | 0.2969 | 0.2009 | 1.00E-39 | 0.1327 | 0.0593 | 5.78E-12 | 0.2725 | 0.1868 | 9.43E-37 |
| S9 | 0.3289 | 0.2446 | 5.36E-43 | 0.2775 | 0.1641 | 1.31E-27 | -0.0130 | 3.28E-4 | 6.38E-01 | 0.3258 | 0.2519 | 2.01E-44 |
| S10 | 0.2136 | 0.0618 | 2.87E-16 | 0.1492 | 0.0277 | 5.86E-08 | -0.0320 | 0.0013 | 2.41E-01 | 0.2145 | 0.0648 | 5.10E-17 |
| S11 | 0.2033 | 0.0509 | 6.62E-13 | 0.1769 | 0.0328 | 9.44E-09 | 0.0555 | 0.0032 | 7.59E-02 | 0.1810 | 0.0424 | 5.78E-11 |
| S12 | 0.2287 | 0.0797 | 2.62E-18 | 0.2434 | 0.0855 | 1.42E-19 | 0.1593 | 0.0351 | 1.06E-08 | 0.1814 | 0.0527 | 1.85E-12 |
| S13 | 0.2697 | 0.0905 | 6.89E-20 | 0.2490 | 0.0735 | 2.60E-16 | 0.1060 | 0.0127 | 8.16E-04 | 0.2467 | 0.0793 | 1.60E-17 |
| S14 | 0.1062 | 0.0153 | 3.11E-04 | 0.0883 | 0.0103 | 3.08E-03 | 0.0462 | 0.0030 | 1.13E-01 | 0.0592 | 0.0049 | 4.19E-02 |
| S15 | 0.2537 | 0.0818 | 6.84E-17 | 0.2710 | 0.0892 | 2.40E-18 | 0.1104 | 0.0159 | 2.99E-04 | 0.2268 | 0.0668 | 5.75E-14 |

**Table S4:** Average regression coefficient of determination (*R*^2^) and Root Mean Squared Error (RMSE). Results are averaged over 1000 bootstraps with 95% confidence intervals for each subject on each model trained. All RMSE values are smaller than the RMSE from a model trained on shuffled labels.

| | $R^2$ | | RMSE | |
| --- | --- | --- | --- | --- |
|  | Subsampled Spectra | Spectral Features | Subsampled Spectra | Spectral Features |
| S1 | 0.203 (0.006) | 0.057 (0.005) | 0.724 (0.003) | 0.780 (0.003) |
| S2 | -0.012 (0.006) | 0.052 (0.005) | 0.822 (0.004) | 0.793 (0.004) |
| S3 | 0.422 (0.006) | 0.237 (0.005) | 0.961 (0.005) | 0.971 (0.004) |
| S4 | 0.173 (0.008) | 0.067 (0.005) | 1.217 (0.006) | 1.095 (0.004) |
| S5 | -0.041 (0.005) | -0.036 (0.004) | 0.914 (0.003) | 0.880 (0.003) |
| S6 | 0.278 (0.005) | 0.242 (0.004) | 0.896 (0.004) | 0.924 (0.004) |
| S7 | 0.326 (0.008) | 0.167 (0.005) | 0.913 (0.005) | 1.015 (0.004) |
| S8 | 0.266 (0.006) | 0.173 (0.004) | 1.112 (0.004) | 1.151 (0.004) |
| S9 | 0.389 (0.004) | 0.342 (0.005) | 1.065 (0.004) | 1.106 (0.004) |
| S10 | 0.253 (0.004) | 0.060 (0.004) | 0.909 (0.003) | 1.022 (0.003) |
| S11 | -0.078 (0.008) | 0.002 (0.003) | 0.787 (0.004) | 0.914 (0.003) |
| S12 | 0.120 (0.006) | 0.049 (0.004) | 0.999 (0.004) | 1.033 (0.003) |
| S13 | 0.147 (0.005) | 0.104 (0.004) | 0.903 (0.003) | 0.928 (0.003) |
| S14 | 0.149 (0.007) | -0.023 (0.003) | 0.790 (0.004) | 0.947 (0.003) |
| S15 | 0.197 (0.005) | 0.044 (0.004) | 0.803 (0.003) | 0.927 (0.003) |
| COMBINED | 0.171 (0.001) | 0.123 (0.001) | 0.949 (0.002) | 0.976 (0.001) |

**Table S5.** Simulated HRF parameters.

|  | TTP (s) | FWHM (s) | Peak PSC<br>(BOLD % Signal Change) |
| --- | --- | --- | --- |
| HRF #1 | 6.0 | 5 | 6.0 |
| HRF #2 | 4.5 | 3.5 | 5.0 |
| HRF #3 | 3.5 | 3.0 | 2.5 |
| HRF #4 | 3.0 | 3.0 | 2.0 |
| HRF #5 | 2.5 | 2.0 | 1.5 |
| HRF #6 | 6.5 | 5.5 | 1.0 |

